# Transverse chromatic offsets with pupil displacements in the human eye: Sources of variability and methods for real-time correction

**DOI:** 10.1101/484386

**Authors:** Alexandra E. Boehm, Claudio M. Privitera, Brian P. Schmidt, Austin Roorda

**Affiliations:** Vision Science Graduate Group, University of California, Berkeley; Berkeley, CA 94720, USA; School of Optometry, University of California, Berkeley; Berkeley, CA 94720, USA

## Abstract

Tracking SLO systems equipped to perform retinally targeted stimulus delivery typically use near-IR wavelengths for retinal imaging and eye tracking and visible wavelengths for stimulation. The lateral offsets between wavelengths caused by transverse chromatic aberration (TCA) must be carefully corrected in order to deliver targeted stimuli to the correct location on the retina. However, both the magnitude and direction of the TCA offset is dependent on the position of the eye’s pupil relative to the incoming beam, and thus can change dynamically within an experimental session without proper control of the pupil position. The goals of this study were twofold: 1) To assess sources of variability in TCA alignments as a function of pupil displacements in an SLO and 2) To demonstrate a novel method for real-time correction of chromatic offsets. To summarize, we found substantial between- and within-subject variability in TCA in the presence of monochromatic aberrations. When adaptive optics was used to fully correct for monochromatic aberrations, variability both within and between observers was minimized. In a second experiment, we demonstrate that pupil tracking can be used to update stimulus delivery in the SLO in real time to correct for variability in chromatic offsets with pupil displacements.

## 1. Introduction

In the last decade, scanning laser ophthalmoscopes [1], or SLOs, both with [2] and without [3] adaptive optics [4], have been used to simultaneously image the retina, track eye motion [5], and deliver targeted, stabilized stimuli to the retina [6,7]. These systems have several advantages for both clinical research and psychophysics. For example, AOSLO microperimetry [8] has been used to assess visual function in local areas of interest in diseased retinas [9]. Additionally, the capability for cone-targeted stimulus delivery with adaptive optics (AO) has made it possible to perform psychophysical experiments at the level of single cone photoreceptors [10-13].

In multi-wavelength tracking SLOs, near-infrared wavelengths are typically used for retinal imaging and eye tracking and visible wavelengths in the 500-700 nm range are used for retinal stimulation. Because the infrared images are used to identify the targeted retinal locations for functional testing, chromatic aberration between the imaging and stimulus wavelengths must be carefully measured and corrected to achieve cellular resolution. Chromatic aberration has two components, longitudinal chromatic aberration (LCA) and transverse chromatic aberration (TCA).

LCA is the difference in axial focus between wavelengths. The magnitude of LCA in human eyes has been carefully quantified [14-19], with estimates of approximately 2 diopters (D) across the visible spectrum, with little variability between individuals.

TCA is the angular offset between the chief rays of two or more wavelengths which causes a lateral offset in the retinal image plane. TCA is typically measured subjectively with a two-wavelength Vernier alignment task [20-23], or objectively with image-based methods in an adaptive optics SLO [24-26]. TCA is highly variable between individuals [17,22,27,28], owing to differences in the position of the achromatic axis in the eye, pupil centration, and alignments of the various optical components of the eye.

Because LCA varies little between individuals [29], a fixed correction between the relative source vergences is sufficient for multi-wavelength imaging and stimulation [24,30]. TCA, however, is a much more challenging problem to account for. While TCA can be measured quickly and reliably with image-based methods in an AOSLO [24], it depends on the exact position of the pupil relative to the incoming beam [23] and thus changes dynamically with small shifts in pupil position.

Previous experiments have accounted for changes in TCA by measuring TCA image offsets at the beginning and end of an experimental session and discarding data from sessions where TCA changed more than a predetermined threshold. However, this ignores any dynamic changes in TCA that occur during an experimental session that could in some cases displace a stimulus from its targeted retinal location by as much as 1 arcmin [25]. Maintaining pupil alignment is especially difficult with unpracticed subjects or when a bite bar is not used, as is often the case with clinical populations and naïve participants. Therefore, a new strategy is needed for measuring and correcting for TCA offsets in real time in order to increase the efficiency of behavioral data collection in an SLO and reduce inter-trial variability in stimulus delivery.

For a perfectly centered optical system, TCA is zero along the optical axis. But in the human eye, TCA will often be present along the line of sight because there is no true optical axis of the human eye and, even if there were, the fovea is displaced relative to it. Consequently, when the subject views a stimulus composed of two wavelengths foveally through a centered pupil, there will be an offset between the two images formed on the retina that is caused by TCA. Displacements from a centered pupil cause the magnitude of the chromatic offset to change, following a linear relationship at small pupil offsets (< 2mm) [20,22,23,25]:

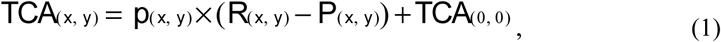

where *TCA_(x, y)_* is the horizontal (*x*) and vertical (*y*) TCA offset in retinal image space in arcminutes, *R*(*_x,y_*) is the (arbitrary) reference pupil position, and *P_(x, y)_* is the real-time pupil position. The pupil offset, *R_(x, y)_* – *P_(x, y)_*, is specified in millimeters. *TCA_(0, 0)_* is the TCA offset at the reference pupil position, and is the y-intercept of the linear function relating TCA to pupil offset. Finally, *p_(x, y)_* is the slope, or the change in TCA offset per unit pupil displacement.

Moving TCA_(0,0)_ to the left hand side of equation 1 yields equation (2), where ΔTCA is the change in TCA offset relative to its value at the reference position.

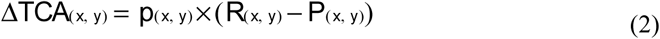

The linear relationship described above implies that changes in TCA with pupil displacement can be easily predicted, and thus compensated for, with only knowledge of the current pupil position relative to some reference and the ΔTCA/mm ratio, or *p_(x,,y)_* in Equations 1 and 2.

The primary goal of the current study was to demonstrate a method for real-time correction of TCA offsets with pupil displacements. However, for TCA compensation in a tracking SLO, it is important to know to what extent variability in *p_(x,,y)_* should be expected. If there is little variability between individuals, the need to calculate slopes for individual subjects will be removed. This would be especially valuable in situations where time with subjects is limited, particularly in studies utilizing clinical populations. Additionally, it is often impractical to measure TCA with image-based cross correlation methods, due to the uncomfortable brightness of the imaging light that is experienced with visible wavelengths and the overall reduction in image quality that is often seen in patient populations.

Previous studies investigating the relationship between pupil displacements and changes in TCA did not directly assess variability between individuals, or within an individual along different pupil meridians. One study [28] reported a correlation between foveal TCA and the magnitude of monochromatic aberrations, though they did not measure TCA with pupil displacements. It remains unknown how monochromatic aberrations might affect TCA.

The current study comprised two experiments. Two systems were used, an adaptive optics scanning laser ophthalmoscope (AOSLO) and a tracking scanning laser ophthalmoscope (TSLO). Because the methods for the two experiments are different, each experiment will be described and discussed separately. In Experiment 1, we measured TCA as a function of vertical and horizontal pupil position in normal eyes in an AOSLO both objectively with image-based methods and subjectively with a Vernier-like alignment technique. Subjective TCA was measured with and without correction for monochromatic aberrations to investigate the sources of variability affecting TCA across the pupil. In Experiment 2, we implemented and tested a strategy to compensate for dynamic changes in TCA in real time during a psychophysics experiment in a TSLO without correction for monochromatic aberrations. To summarize, we found that, in the presence of aberration, there are significant differences in the slope of TCA with pupil offsets between meridians in the eye and between individuals. After correcting aberrations, the measurements are much less variable. Finally, by performing a calibration of the slope of TCA with pupil offsets, we can use pupil tracking to guide compensation of TCA offsets in real time.

## 2. Experiment 1: Sources of variability in TCA with pupil displacements

### 2.1 Subjects

Two subjects (20075 and 20076, ages 29 and 30) participated in Experiment 1. Both were authors of the study. All procedures were approved by the University of California, Berkeley Committee for Protection of Human Subjects and adhered to the tenets of the Declaration of Helsinki. All subjects signed informed consent documents before participating in the experiments.

### 2.2 Apparatus

A multi-wavelength adaptive optics scanning laser ophthalmoscope (AOSLO) was used for imaging and stimulus delivery. This system is described in detail elsewhere [2,10], with relevant details specified below.

Three wavelengths were selected from a supercontinuum light source (SuperK EXTREME, NKT Photonics, Birkerod, Denmark) and the vergence of each channel was adjusted to compensate for LCA of the human eye [20]. 940 nm light was used for wavefront sensing. 840 nm (near-IR) and 543 nm (green) light were used for both imaging and stimulation. Beam size at the pupil was approximately 3 mm in the subjective alignment task, and 6.5 mm for retinal imaging. All three channels were raster-scanned onto the retina and subtended a square field 0.95 x 0.95 degrees in size (corresponding to a sampling resolution of 0.11 arcmin per pixel). Light backscattered from the 840 and 543 nm channels were collected into a photomultiplier tube (Photosensor module H7422, Hamamatsu Photonics, Hamamatsu, Japan) via a confocal pinhole and digitized. The intensity of the 840 and 543 nm channels were modulated with fiber-coupled acousto-optic modulators (Brimrose Corp, Sparks Glencoe, MD). The subject’s head was positioned with a bite bar mounted on an X,Y,Z translation stage.

Monochromatic aberrations were corrected with a deformable mirror (DM97, ALPAO, Montbonnot-Saint-Martin, France) operated in one of three modes, described in further detail in *2.4 Procedure*.

Pupil videos were recorded using a 1/2 inch USB CMOS color board camera (DFM 61BUC02-ML, ImagingSource Inc. Charlotte, NC) at 1024 x 768 pixel resolution corresponding to approximately 60 pixels per mm. The sensor was controlled by a MATLAB plugin which allows full control and optimization of the camera’s settings (e.g. exposure, gain, gamma) from the MATLAB working space. Images of the pupil were formed via retro-illumination from the 840 nm imaging beam (i.e. red-eye reflex). Videos were acquired directly in MATLAB at approximately 10 fps and then converted into gray scale. The shape of the pupil was first roughly separated by applying a simple gray scale threshold and then its contour was finely defined by a gradient (edge detection) operator. Next, an ellipse was fitted using a least-squares criterion. Pupil tracking was represented by the x and y coordinates of the center of the fitted ellipse.

The pupil camera was coaligned with the AOSLO beam path by means of a beamsplitter placed before the eye with a 45° angle of incidence and broad-spectrum anti-reflective coating on one side to minimize reflections (Ross Optical Industries, El Paso, Texas). The pupil camera was placed at the back end of a Maxwellian assembly which is typically used to form background images and fixation targets, though the projector was not used in the current study. A 50 mm square hot mirror (#43-452, Edmund Optics, Barrington, NJ) was placed before the projector to reflect light to the pupil camera while allowing transmittance of visible wavelengths from the projector. The optical axes of the pupil camera and SLO beam path were carefully aligned to ensure that axial misalignments of the subject’s pupil were not misconstrued as lateral displacements.

### 2.3 Stimulus

The stimulus consisted of four dark squares, 4.42 arcminutes in length, arranged in a square pattern, and a green target square (λ = 543 nm) of the same size that was spatially offset by a random amount between 0 and 2.21 arcminutes (vertically and horizontally) of the objective center of the stimulus configuration at the beginning of each trial (Fig. 1A). The decrements were achieved by switching off the 840 nm wavelength at the appropriate points in the raster scan. The background was a square raster pattern subtending 0.954 deg and was composed of the 840 nm light source that is typically used for imaging in addition to leak from the 543 nm light source due to first order light scatter through the AOM [31].

**Fig 1:**
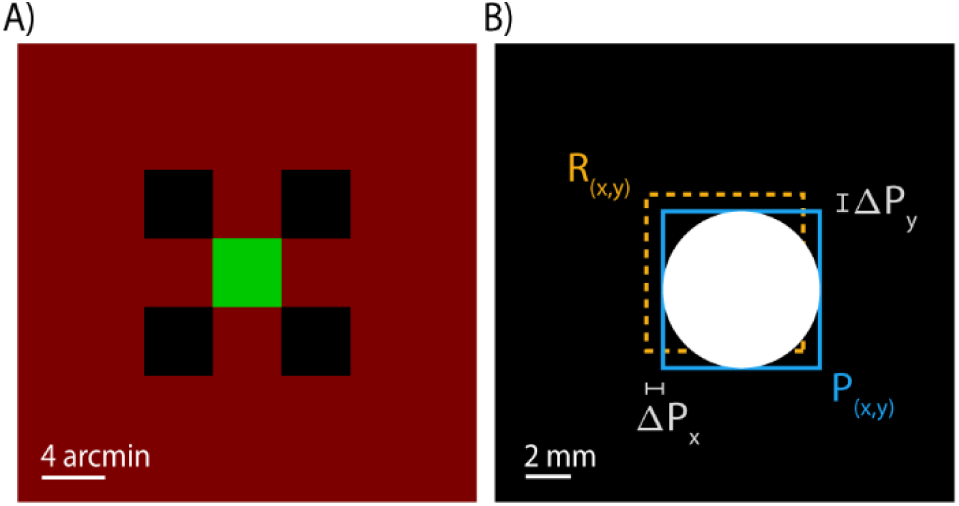
TCA induced by pupil displacements. (A) The stimulus used for the alignment task. Subjects aligned a green (λ = 543 nm) target square to be centered with respect to the surrounding decrement (λ = 840 nm) squares. (B) Pupil tracking was performed in real time throughout the experiment. The experimenter induced pupil offsets relative to the reference pupil position, *R_(x,y)_*.

### 2.4 Procedure

Subjects’ pupils were dilated and cyclopleged with 1-2 drops of 1% tropicamide ophthalmic solution 15 minutes prior to the experiment. The experimenter aligned the subject’s pupil to be centered with respect to the imaging beam and this was set as the reference pupil position.

At the beginning of each trial, the experimenter moved the subject’s pupil within +/- 0.5 mm of the reference pupil position in one of nine positions, upward-left, left, downward-left, downward, downward-right, right, upward right, upward, and centered (Fig 1B). The induced pupil offsets followed either a clockwise or counter-clockwise pattern which was alternated after the centered pupil position was reached at the end of each set of trials.

The subject adjusted the position of the green target square using a keyboard until it appeared to be centered with respect to the four decrement squares. Subjects indicated the alignment by pressing the enter key, and pupil positions were recorded at the time of the keypress. Subjects completed 45 alignments in each block of trials, where in each block one of the three modes of optical correction was applied (‘defocus only,’ ‘defocus + astigmatism’, or ‘AO closed-loop’). In the first mode (‘defocus only’), defocus was manually added to the mirror, which was otherwise static for the duration of the session. The point of subjective best-focus was found by scanning the stimulus onto the retina and adjusting defocus until the subject reported seeing the sharpest image. In the second mode (‘defocus + astigmatism’), defocus and astigmatism were manually added to the mirror and the point of subjective best-focus was again found. In the third mode (‘AO closed-loop’), real-time wavefront measurements were used to control the deformable mirror and correct both low and higher-order monochromatic aberrations. In this mode, correction continued in real-time throughout the duration of the session.

Additionally, we measured TCA at similar pupil offsets with image-based methods following the methodology described in detail in Harmening et al. [24]. Briefly, AOSLO images were recorded with λ = 543 nm (4.5 µW) and λ = 843 nm (52 µW) light simultaneously, by interleaving lines composed of light from the different channels. A cross-correlation procedure was used to calculate the image offset between the two wavelengths. Note that the image-based methods were only possible in the AO closed-loop mode – the low signal and poor resolution precluded image-based alignments for the other modes.

### 2.5 Results

Results for the two subjects’ alignments under the three optical correction conditions and objective measurements of foveal TCA are shown in Fig. 2. In the figure, each point represents the vertical or horizontal offset the subject required to align the 543 nm increment square to the 840 nm decrement squares, plotted as a function of pupil offset in the respective direction. Each subject’s data were fit to Eq. 2, where the slopes (*p_(x,y)_*) represent the change in TCA per unit pupil displacement.

**Fig 2:**
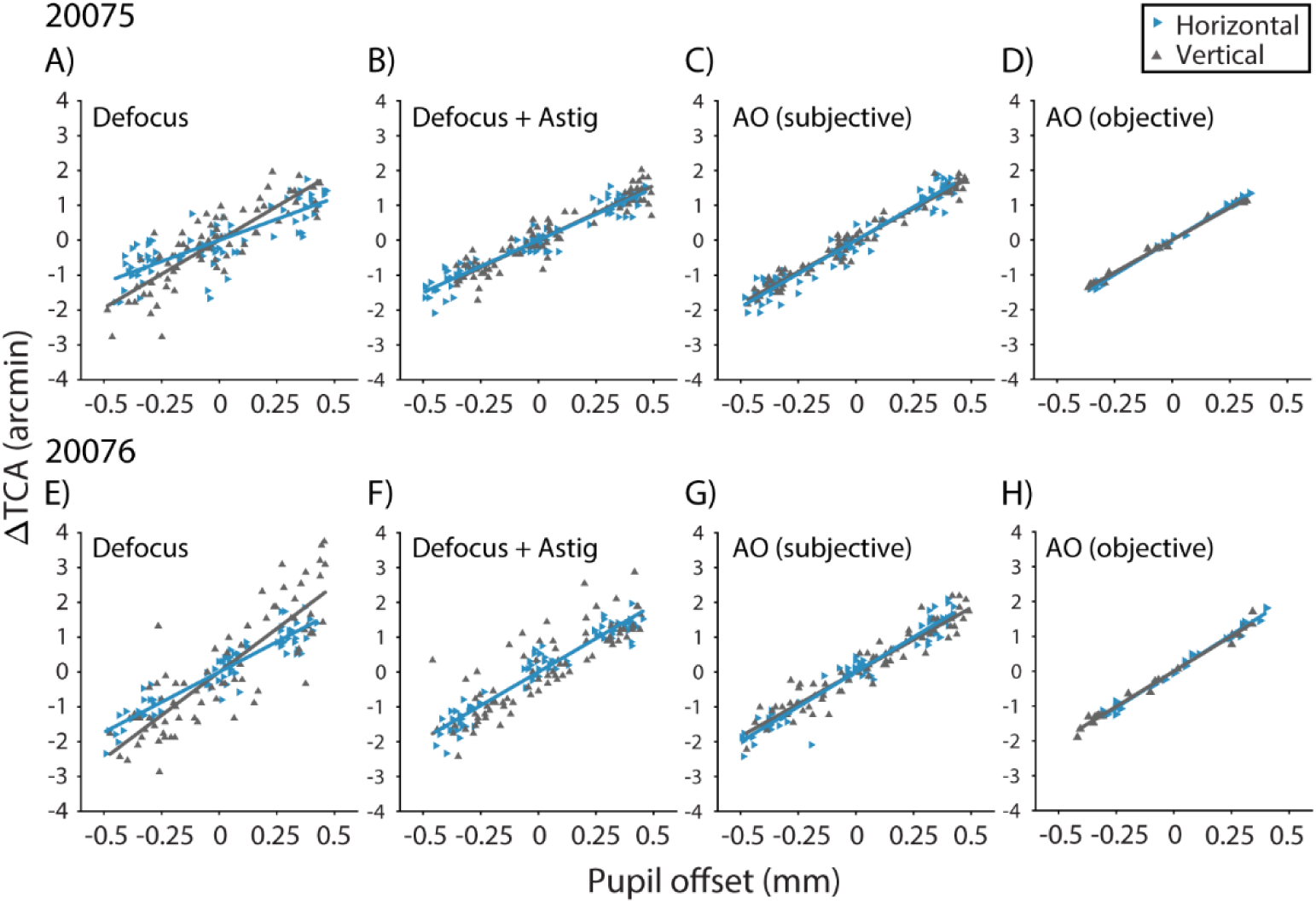
Changes in TCA (λ = 543 and 840 nm) with pupil displacement in the AOSLO for subjects 20075 (A-D) and 20076 (E-H). TCA was measured subjectively under different optical corrections: Defocus only (panels A and E), defocus and astigmatism (B and F), and AO (C and G), and objectively with AO image-based methods (D and H). The horizontal (blue rightward triangles) and vertical (gray upward triangles) components of the chromatic offset follow a linear relationship with the magnitude of the horizontal and vertical pupil offsets, respectively.

Bar charts showing the slope estimates and statistical comparisons are shown in Fig 3. Slopes of the regression lines were compared using the z-test method explained by Kleinbaum et al. [32], with z-scores evaluated as the difference between regression slopes divided by the

**Fig 3:**
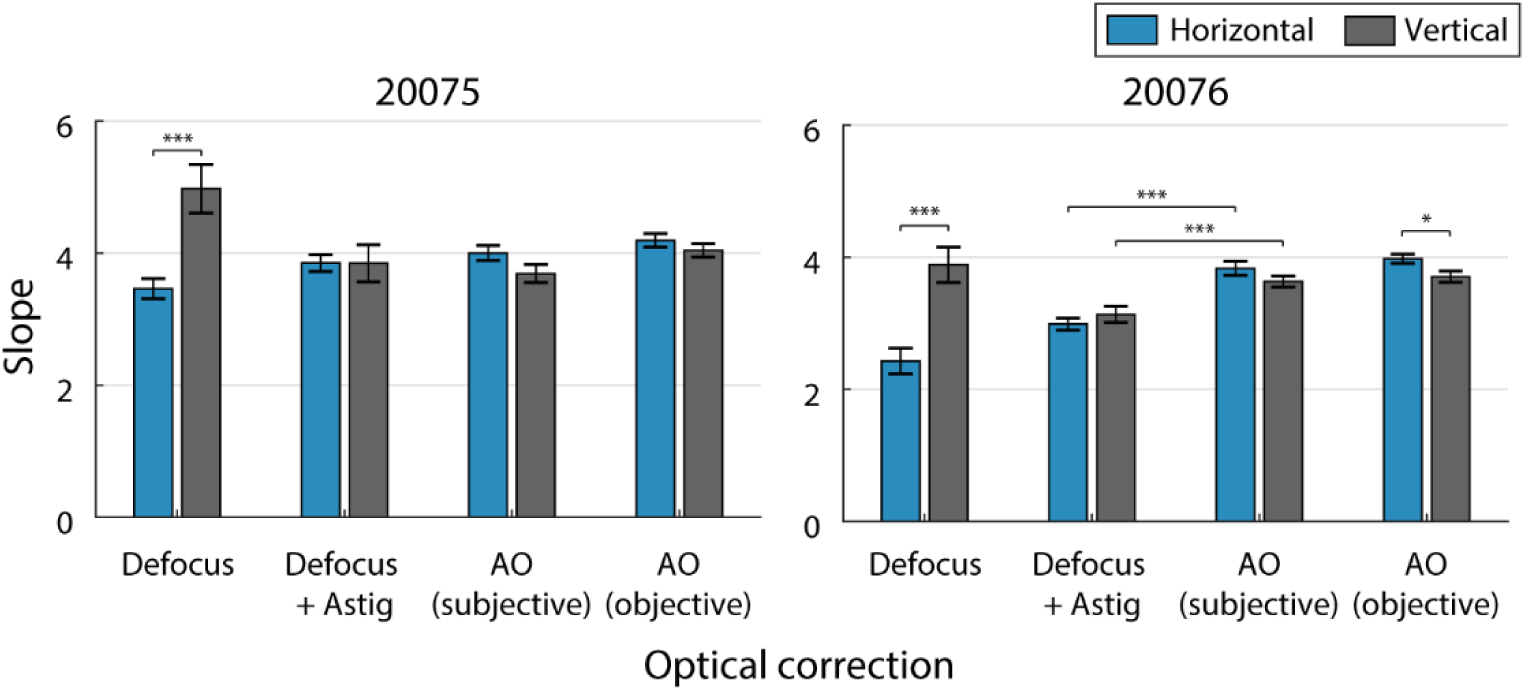
Slopes for horizontal (blue) and vertical (gray) changes in TCA with pupil displacements for subjects 20075 (left) and 20076 (right) under different optical corrections: defocus, defocus + astigmatism, and AO. Slope estimates for changes in TCA with AO correction are reported for two measurement methods, subjective alignments and objective image-based. Error bars are standard errors of the slope estimates. Brackets show statistically significant comparisons (* p < 0.05, ** p < 0.01, *** p < 0.001).

In the defocus correction condition (Fig. 2A and E), slopes relating the horizontal component of TCA to horizontal pupil offsets were significantly different from vertical changes in TCA with vertical pupil offsets (*p* < 10^−4^; Fig. 3). However, when we applied a correction for astigmatism (Fig. 2B and F), the differences between horizontal and vertical slopes were no longer significant. This was also the case with AO correction (Fig. 2C and G); vertical and horizontal slopes were not significantly different for either subject (Fig. 3).

Next, the amount of variability in TCA captured by the linear model was quantified with a correlation analysis (R^2^). A perfectly linear system with zero noise would have R^2^ = 1. In both subjects, R^2^ increased as monochromatic aberrations were reduced. For 20075, R^2^ increased from 0.7993 (average of horizontal and vertical R^2^) when only defocus was corrected to 0.8195 when astigmatism was corrected as well. R^2^ increased further to 0.9270 when AO correction was applied. For 20076, R^2^ was 0.7020 for defocus only, 0.9069 for defocus + astigmatism, and 0.9524 with AO correction. Thus, as monochromatic blur was removed, variability in subjective TCA decreased.

Additionally, we compared the slopes between the astigmatism and AO correction conditions to see if there was any further change in slope after correction for higher order aberrations. For 20075, slopes were not significantly different between the two conditions. However, both *p_x_* and *p_y_* were slightly steeper in the AO correction condition for 20076 (*p* < 10^−4^). Comparing the horizontal and vertical slopes across the two subjects, we found that they were not significantly different in the AO correction condition, with a mean *p* ratio of 3.845 for 20075 and 3.731 for 20076.

Also under the AO correction condition, we measured foveal TCA with pupil displacements using an image-based cross correlation procedure (Fig. 2D and H). In this case, there was a small difference between the horizontal and vertical slopes for 20076 only (*p* < 0.05), but not for 20075 (Fig. 3). The average slope for 20075 was 4.117 (SEM = 0.0756). Additionally, there was a small but statistically significant difference in slopes between the two subjects for vertical TCA with vertical pupil offsets only (*p* < 0.05). The differences in TCA measured with image-based methods after AO correction could be due to differences in corneal curvature between subjects and/or between pupil meridians. We discuss this point further below.

Finally, we compared the slopes of the TCA versus pupil offset functions between the subjective alignment task to those acquired with the image-based method. For 20076, there was no significant difference between the slopes calculated with the two different methodologies. For 20075, there was a small difference in the vertical TCA with vertical pupil displacements, but this difference was not statistically significant at the α = 0.05 level after correction for multiple comparisons (Benjamini-Hochberg procedure [33]).

### 2.6 Discussion

In Experiment 1, we compared slopes of two subject’s TCA versus pupil offset functions in an AOSLO under different optical corrections: defocus only, defocus + astigmatism, and AO. We found that the differences between slopes with respect to pupil meridian were removed when subjects made alignments with astigmatism correction and AO. We first explored whether the difference in slope could be caused by meridional differences in corneal curvature. But when we varied the radius of curvature in a simple chromatic eye model [20] it did not produce differences between *p_x_* and *p_y_* as large as we observed in our data, suggesting that meridional differences in corneal curvature cannot fully account for this. Varying the radius of curvature to simulate asymmetries in normal eyes [34], however, did produce differences in TCA consistent with the small differences observed with AO image-based methods, suggesting that small differences in TCA (< 0.5 arcmin) present even after AO correction may be due to differences in corneal curvature.

We then considered how uncorrected monochromatic aberrations might affect TCA. One possibility is that changes in monochromatic aberrations with wavelength could add additional perceived offsets between polychromatic targets, since even monochromatic targets appear misaligned by an amount proportional to the aberration when viewed through split-field systems where one target is presented in normal view and the other through an isolated portion of the pupil [35]. However, Marcos et al. [17] found variations in monochromatic aberrations with wavelength to be very small, and therefore they are unlikely to result in the relatively large differences in slope that we observed in our subjects.

Marcos et al. [28] investigated variability in foveal TCA, and concluded that the individual differences in foveal TCA could not be fully explained by the displacement of the achromatic axis from the optical axis of the eye, pupil centration, corneal curvature or asphericity, or alignment of the various optical components of the eye. Instead, they noted a correlation between the magnitude of monochromatic aberrations (root mean square, or RMS) and foveal TCA which they attributed to irregularities in the shape of the cornea and lens. However, given that changes in corneal curvature have small effects on the magnitude of TCA, it is not clear how monochromatic aberrations could generate the differences they observed.

Another possibility is that the differences in actual TCA are very small in the eyes we tested, and the observed differences reflect an observer bias in perceived offsets between the targets that is due to overall blur of the stimulus [36-39]. Moreover, as blur is expected to change with increasing offset from a central pupil position, these biases will not only give rise to variability in alignments but differences in slope as well. Fig. 4 offers a visual to help explain this effect. An aligned stimulus was convolved with the computed point spread, shown in panel B, of a typical eye over a 3 mm pupil that was displaced laterally in the horizontal and vertical directions (Fig. 4A). As can be seen, the blur from the PSF makes the correct alignment ambiguous and can change the subjective alignment depending on the offset position, thereby affecting the slope. We also suspect that the variability in perceived alignment will be further exacerbated by the reverse polarity of the features that are being aligned [39-44].

**Fig 4:**
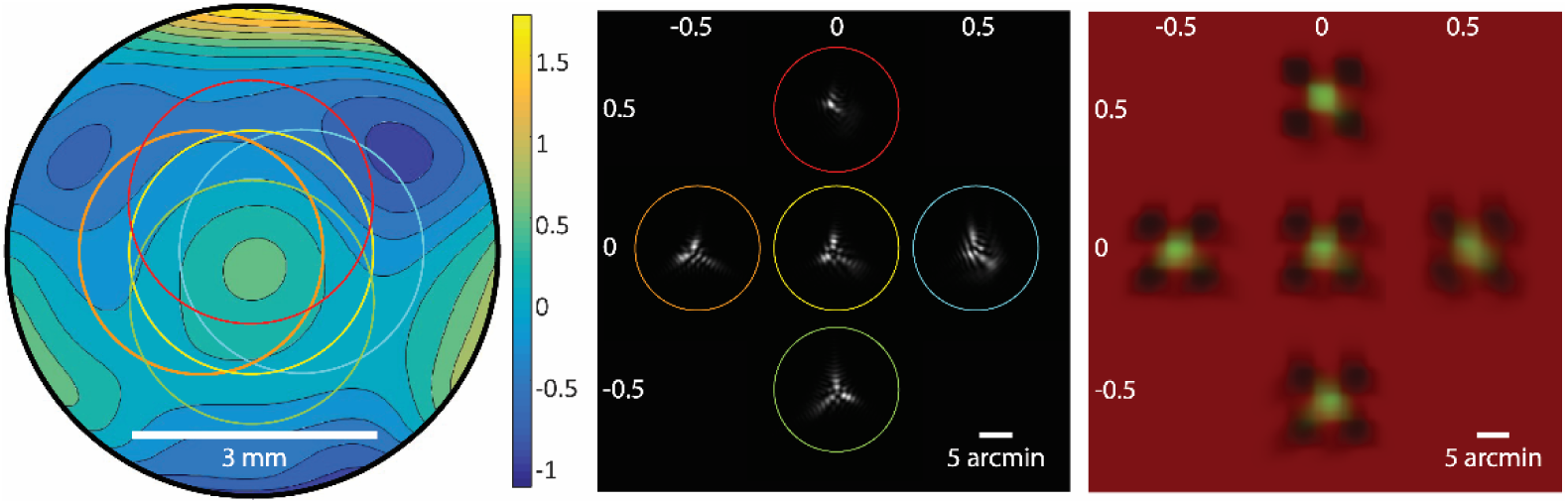
Simulations of the appearance of the TCA stimulus as a function of pupil position. A) Model wavefront of a typical eye used for simulations. Units for the color bar are micrometers. Colored circles overlaid on the wavefront image demarcate the area of the pupil which was sampled to generate the PSFs shown in B). Defocus was set to zero for the centered position only (yellow circle). The pupil offsets for each sample relative to the centered position are given in millimeters (mm) along the margins of panels B) and C). C) Simulations of the appearance of the stimulus, generated by convolving the PSFs shown in B) with the aligned stimulus (Fig. 1A). Each image represents the stimulus appearance at a different vertical or horizontal pupil offset (units are mm). Note that this does not include the offset between stimuli incurred by chromatic aberration, and thus is representative only of the appearance with monochromatic blur.

To our knowledge, we are the first to report significant differences in TCA with pupil position in different meridians. Previous studies using Vernier stimuli viewed through a pinhole aperture only measured the horizontal component of the TCA offset as a function of pupil displacement [20,22,23]. These authors did not directly assess variability between subjects, however their data demonstrate individual differences in slopes for horizontal TCA comparable to those observed here. Privitera et al. [25] measured TCA offsets for both horizontal and vertical pupil displacements with AOSLO image-based methods, and the small difference between the horizontal and vertical slopes in their subject is consistent with the small variability observed in the AO imaging results for our two subjects.

We found an average *p* ratio of 3.98 with image-based methods, slightly higher than what was reported previously by our lab [25]. This means that a pupil misalignment of only 0.13 mm from the reference position (where TCA is measured at the beginning of an experiment with image-based methods) will displace a stimulus from its targeted retinal location by half an arcmin. This has implications for single-cone targeted stimulus delivery where authors are interested in relating behavioral phenomena to single-cone stimulation. Cone photoreceptor spacing at 1-2 degrees eccentricity, where these experiments are typically performed, is approximately 1 arcmin, leaving little room for error induced by pupil displacements. To combat this, we have introduced real-time visual and auditory feedback from the pupil tracking plugin to assist the subject in keeping their pupil aligned. We find that in many cases this is sufficient and there is no need for active compensation. However, work done closer to the fovea will demand a higher level of stimulus placement accuracy and a better strategy for TCA compensation, like we demonstrate in Experiment 2. Additionally, TCA compensation would be useful for naïve subjects and particularly clinical populations who may have difficulty maintaining alignment.

In Experiment 2, we used *p* ratios measured for individual subjects in order to compensate for individual differences in TCA offset with pupil displacements. We used a TSLO to demonstrate TCA compensation without correction for monochromatic aberrations, other than defocus. Implementation in an AOSLO would be the same.

## 3. Experiment 2: Real-time correction for changes in TCA with pupil displacements

### 3.1 Subjects

Eight subjects (4 males, 4 females ages 23-50), none of whom participated in the previous experiment, participated in Experiment 2. Subjects 10003 and 20036 were authors of this study, the remaining six subjects were naïve to the purposes of the experiment.

### 3.2 Apparatus

The TSLO used in these experiments is described elsewhere [3], with details specific to these experiments described briefly here. The infrared light source was an 840 nm superluminescent diode (Superlum, Moscow, Russia). A second light source, a diode-pumped solid-state laser (CrystaLaser, Reno, NV) with λ = 532 nm, was aligned so that its beam path was coaligned with the infrared wavelength at a dichroic beam splitter placed before the entrance pupil. Two fiber-coupled acousto-optic modulators (Brimrose Corp, Sparks Glencoe, MD) were used to independently modulate the intensities of the light sources. A chin rest and temple pads were mounted on an X,Y,Z stage that the experimenter used to align the subject’s pupil to the pupil plane of the system.

Pupil tracking in the TSLO system was the same as described in Experiment 1.

### 3.3 Stimulus

The stimulus configuration was the same as in Experiment 1 (Fig. 1A), except that the background composed of the 840 nm raster was 3.2 deg (reflecting the larger field of view in the TSLO system) and each square was 11.25 arcminutes in length. Additionally, the target square was composed of 532 nm light.

### 3.4 Procedure

Subjects’ pupils were dilated and cyclopleged with 1-2 drops of 1% tropicamide ophthalmic solution 15 minutes prior to the experiment. A trial lens was placed at the entrance pupil of the TSLO to correct for defocus when necessary. The experimenter aligned the subject’s pupil to be centered with respect to the imaging beam and this was set as the reference pupil position.

Subjects completed 180 trials in each experimental session. Each trial was the same as described in Experiment 1 (see *2.4 Procedure*), except that the pupil offsets extended to +/- 0.75 mm of the reference pupil position. Three subjects (20093, 20106, 10003) completed one session and then went on to participate in the compensation task, described in the following paragraph, while the other five subjects completed two sessions without compensation to further assess individual differences in slopes.

To compensate for changes in TCA with pupil displacements, the position of the green (λ = 532 nm) increment square generated by the stimulus channel was updated on each frame by an amount equal and opposite to the expected TCA offset, inferred from the subject’s current pupil position and their previously determined *p* ratio. This was achieved by real-time feedback from the pupil tracker which was used to estimate the TCA offset and then update the timing of the green stimulus delivery in the raster scan accordingly. Subjects used a keyboard to align the increment square to be centered relative to the decrement squares. Spatial offsets indicated by the subject were summed with any additional offsets intended to compensate for changes in TCA.

Subjects completed 90 trials where compensation was or was not applied. In the no-compensation condition, the position of the increment pattern was updated by subject’s key presses only, and thus was no different from the calibration procedure. Trials were pseudo-randomly interleaved, with 45 in each condition.

### 3.5 Results

Fig. 5 shows data and fits to Equation 2 for three subjects. Results for the remaining five subjects were similar, with differences between subjects described in detail below and summarized in Fig. 6. In Fig. 5, each data point represents the vertical or horizontal offset the subject required to align the 532 nm target square to the 840 nm decrement pattern, plotted as a function of pupil offset in the respective direction. Each subject’s data were fit to Eq. 2, where the slopes (*p_(x,y)_*) represent the change in TCA per unit pupil displacement.

**Fig 5:**
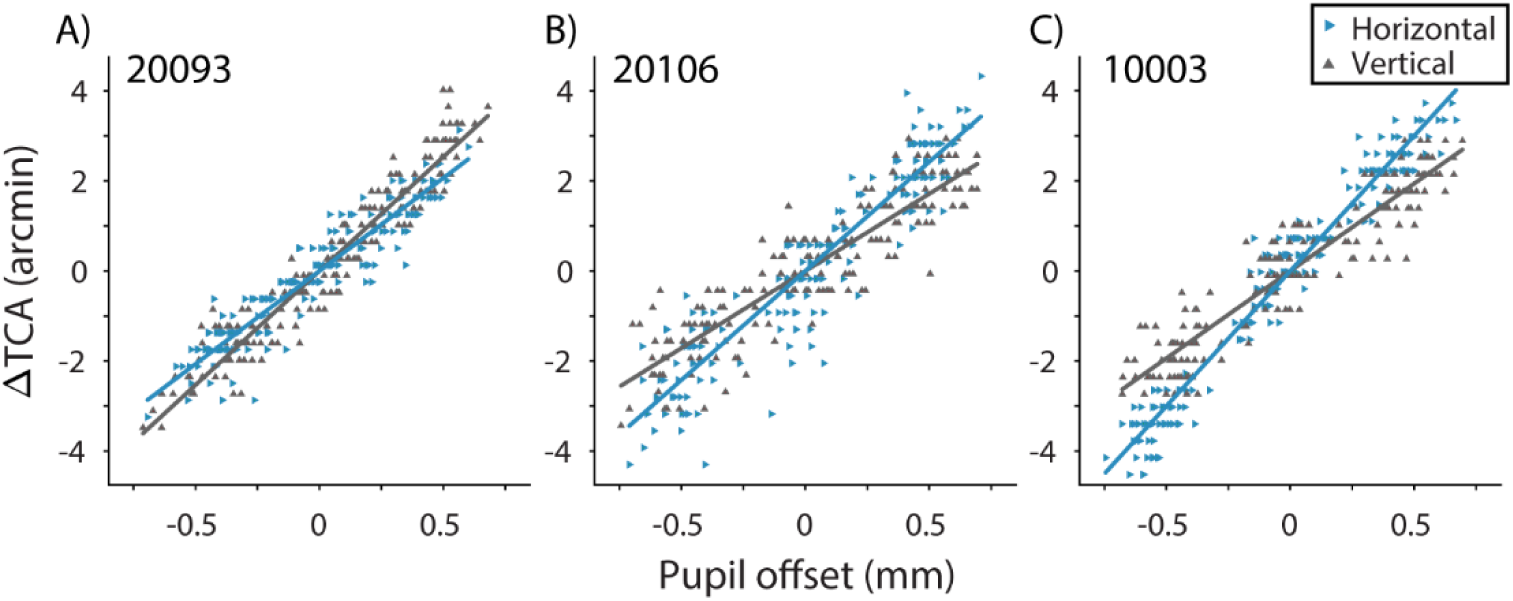
Subjective alignments of the stimulus at different pupil positions for three subjects: A) 20093, B) 20106, and C) 10003. Pupil offsets are relative to the reference pupil position at zero. Subjects’ alignments are expressed as ΔTCA, the change in TCA offset (λ = 532 and 840 nm) relative to its magnitude at the reference pupil position. Data were fit with ordinary least squares (OLS) regression separately for horizontal (blue rightward triangles) and vertical (gray upward triangles) changes in TCA for horizontal and vertical pupil offsets, respectively.

**Fig 6:**
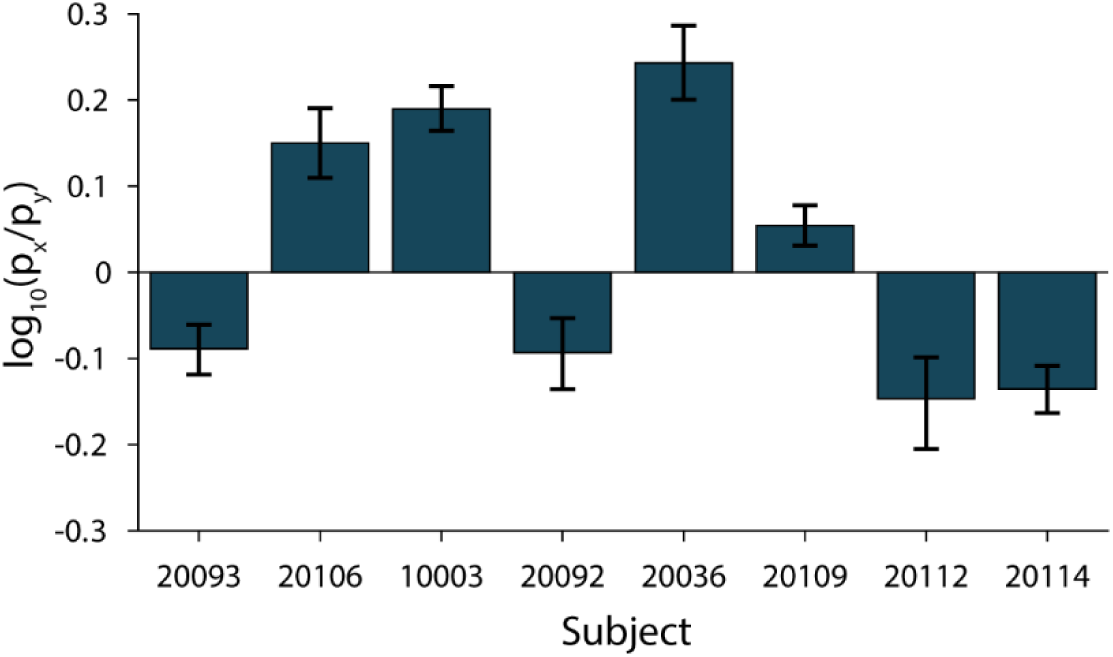
Individual differences in anisotropy of *p_x_* and *p_y_*. The log ratio log_10_(*p_x_*/*p_y_*), is shown for each subject. When *p_x_*= *p_y_*, the log ratio is 0. Points above the line indicate *p_x_* > *p_y_*, and below the line *p_x_* < *p_y_*. Error bars are 95% bootstrapped confidence intervals.

In all 8 subjects, the difference in slopes was significantly different between the two pupil meridians. Fig. 6 shows the log ratios, log_10_(*p_x_*/*p_y_*), for all subjects. When *p_x_* = *p_y_*, the log ratio is zero (solid line). Additionally, there was between-subject variability in both the magnitude and direction of the anisotropy. In half of our subjects *p_x_* > *p_y_* and the other half had *p_x_* < *p_y_*.

Data and fits for the compensation condition are shown in Fig. 7. Following compensation for changes in TCA with pupil displacements, *p_x_* and *p_y_* were not significantly different from zero, except for *p_x_* (the horizontal component of TCA with horizontal pupil displacements) in 20106 (Fig. 7B), which was due to a noisy calibration measurement. Additionally, the within-subject anisotropies in ΔTCA with respect to pupil meridian (Fig. 6) were no longer observed.

**Fig 7:**
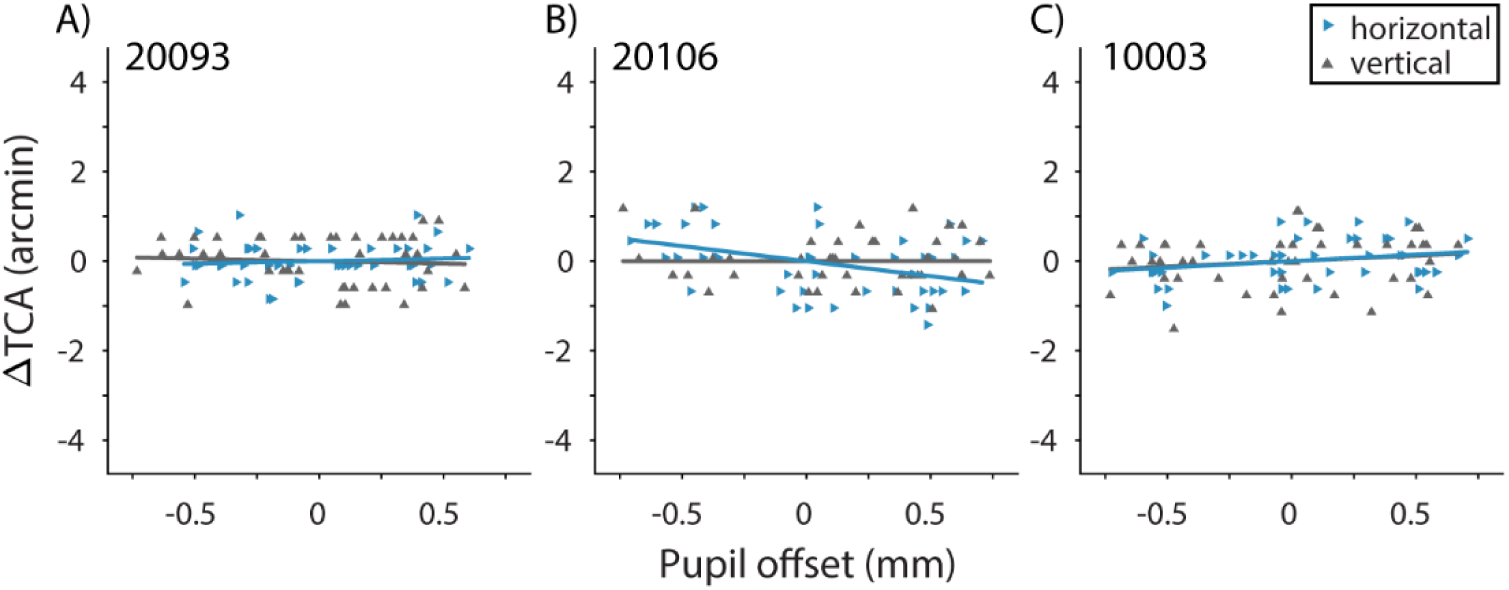
Compensation for ΔTCA with pupil displacement. Results are shown for the same subjects as in Fig. 5. Slopes (*p_(x,y)_*) for horizontal (blue rightward triangles) and vertical (gray upward triangles) pupil offsets were not significantly different from zero, except for *p_x_* in 20106.

We also compared the slopes across the no-compensation condition and the initial estimate determined in the calibration procedure. We found that slopes between the two sessions were not statistically different (*p* > 0.2), except in the case of *p_x_* in 20106. This suggests that our failure to fully compensate for changes in TCA in this case was likely due to an overestimation of *p_x_* in the calibration procedure.

### 3.6 Discussion

In Experiment 2, we used *p* ratios measured for individual subjects in the TSLO in order to compensate for individual differences in TCA offset with pupil displacements without correction for monochromatic aberrations other than defocus. Similar to our results from Experiment 1, we found significant within- and between-subject variability in the slopes of the TCA versus pupil offset functions in all eight subjects.

We used *p* ratios calibrated for individual subjects and both pupil meridians in order to compensate for changes in perceived chromatic offsets with pupil displacements in the TSLO. Perceived changes in TCA with pupil displacements were compensated by a combination of real-time pupil tracking and updating of stimulus delivery. With real-time compensation, slopes of the ΔTCA versus pupil offset function were flattened and there was no longer a systematic relationship between pupil offset and subjects’ alignments, except in the case of the horizontal component of TCA for 20106 which was likely due to an overestimation of *p_x_* in the calibration procedure (see *3.5 Results*). Even in this case, most of the variability in psychophysical alignment due to pupil displacements was removed.

Though we used a TSLO system to demonstrate TCA compensation, implementation in the AOSLO would be effectively the same because both systems are equipped with the same hardware and software components utilized in this technique. However, in AO systems, which correct for monochromatic aberrations across the fully dilated pupil, our results from Experiment 1 suggest it may be possible to use a single *p* ratio for both meridians. Additionally, there was little variability between the two individuals in the TCA versus pupil offset function. The small differences between subjects’ vertical slopes (0.34 arcmin/mm) we observed following AO correction are negligible in the context of TCA compensation, given that foveal cone spacing is approximately 0.5 arcmin [45] and the pupil is unlikely to become displaced by more than a fraction of a millimeter. Overall, these data suggest that a uniform *p* ratio based on the average of a few subjects can be used in systems which utilize AO to correct for higher order aberrations, though a complete characterization of between subject variability will require a larger study.

## 4. Conclusions

In this study we demonstrated that pupil tracking can be used to predict dynamic changes in TCA that occur with small shifts in pupil position. By updating stimulus delivery in accordance with the subjects’ pupil position, we successfully compensated for changes in TCA that occurred with pupil offsets (Experiment 2). We found that changes in TCA followed a linear relationship with pupil offset, consistent with previous estimates based on psychophysics [20,22,23] and AOSLO multi-wavelength image analyses [25]. However, in both experiments we found that there was substantial variability in the slopes of the ΔTCA versus pupil offset functions, both between and within individuals, when monochromatic aberrations were uncorrected, and we propose that much of this variability is due to biases in alignments in the presence of optical blur. In two subjects, these differences were minimized when AO was applied, suggesting that in many cases a uniform slope can be used to correct for TCA across subjects in an AOSLO.

## Acknowledgements

We thank Pavan Tiruveedhula for his technical support.

